# Natural selection reduces linked neutral divergence between distantly related species

**DOI:** 10.1101/031740

**Authors:** Tanya N. Phung, Christian D. Huber, Kirk E. Lohmueller

## Abstract

Much work has been devoted to understanding the evolutionary processes shaping genetic variation across genomes. Studies have found neutral polymorphism is reduced close to genes and in regions of low recombination, suggesting the effects of natural selection. However, the effect of selection on neutral sequence divergence between species remains ambiguous. While studies have reported correlations between divergence and recombination, theoretical arguments suggest selection may not affect divergence at linked neutral sites. Here we address these outstanding issues by examining how natural selection has affected divergence between distantly related species. We show that neutral divergence is negatively correlated with functional content and positively correlated with estimates of background selection from primates. These patterns persist even when comparing humans and mice, species that split 75 million years ago. Further, neutral divergence is positively correlated with recombination rate. The correlation increases when focusing on genic regions, and biased gene conversion cannot explain all of this correlation. These signatures suggest that natural selection has affected linked divergence between distantly related species. Coalescent models indicate that background selection can generate these patterns. Even when the contribution of ancestral polymorphism to divergence is small, background selection in the ancestral population can still explain a large proportion of the variance in divergence across the genome. Thus, the view that selection does not affect divergence at linked neutral sites needs to be reconsidered. Our work has important implications for understanding evolution of genomes and interpreting patterns of genetic variation.

## INTRODUCTION

Determining the evolutionary forces affecting genetic variation has been a central goal in population genetics over the past several decades. A large body of empirical and theoretical work has suggested that neutral genetic variation within a species can be influenced by nearby genetic variants that are affected by natural selection (reviewed in Cutter and Payseur 2013). This can occur via two mechanisms. In a selective sweep, a neutral allele linked to a beneficial mutation will reach high frequency (Maynard Smith and Haigh 1974; Kaplan et al. 1989). Selective sweeps reduce neutral genetic variation surrounding regions of the genome where selection more commonly occurred. The second process, background selection, also reduces neutral genetic variation (Charlesworth et al. 1993; Hudson and Kaplan 1995; Nordborg et al. 1996; Charlesworth 2012a). Here deleterious mutations that are eliminated by purifying selection also remove nearby neutral genetic variation. Many empirical studies have found strong evidence for the effects of background selection and selective sweeps affecting patterns of neutral genetic diversity across the human genome. For example, several studies have reported a correlation between genetic variation and recombination rate (Nachman 2001; Hellmann et al. 2003, 2005, 2008; Cai et al. 2009; Lohmueller et al. 2011). This correlation can be driven by selective sweeps and background selection because these processes affect a larger number of base pairs in areas of the genome with low recombination rate than with high recombination rate. Additionally, other studies found reduced neutral genetic diversity surrounding genes (Payseur and Nachman 2002; Cai et al. 2009; McVicker et al. 2009; Hernandez et al. 2011; Lohmueller et al. 2011; Enard et al. 2014), which is consistent with the idea that there is more selection occurring near functional elements of the genome.

While the evidence for selection reducing genetic diversity is unequivocal, the effect of selection on sequence divergence between species is less clear. Elegant theoretical arguments have suggested selection does not affect the substitution rate at linked neutral sites (Birky and Walsh 1988). However, these theoretical arguments do not include mutations that arose in the common ancestral population. Such ancestral polymorphism is widespread and has been shown to be a significant confounder in estimating population divergence times (Edwards and Beerli 2000). When also including ancestral polymorphisms, it becomes less clear whether selection affects divergence at linked neutral sites. Based on simple coalescent arguments, neutral polymorphism in the ancestral population will be affected by linkage to selected sites the same way as genetic diversity within a population (**Figure 1**). Presumably, neutral divergence between closely related species, with lots of ancestral polymorphism, then could be affected by selection. But, selection is thought to not have an effect on linked neutral divergence when considering species with very long divergence times (Birky and Walsh 1988; Hellmann et al. 2003; Cruickshank and Hahn 2014). The argument for the lack of an effect with long split times is that there would be many opportunities for mutations to occur after the two lineages split (**Figure 1**). These neutral mutations that occur after the split would not be influenced by selection at linked neutral sites (Birky and Walsh 1988) and would dilute the signal from the ancestral polymorphism. Thus, it is generally believed that selection at linked neutral sites should not affect divergence between distantly related species. An example of this argument was presented by Hellmann et al. (2003). They argued that the positive correlation between human-baboon divergence and human recombination was due to mutagenic recombination, rather than selection affecting linked neutral sites, because of the long split time between humans and baboons (>20 million years). Reed et al. (2005) suggested that though it is unlikely background selection by itself could explain the entire correlation observed by Hellman et al., background selection may still contribute to divergence. However, beyond these verbal arguments, there has been little quantitative investigation of the effect that selection has on divergence at linked neutral sites among distantly divergent species when including ancestral polymorphism.

**Figure 1:**
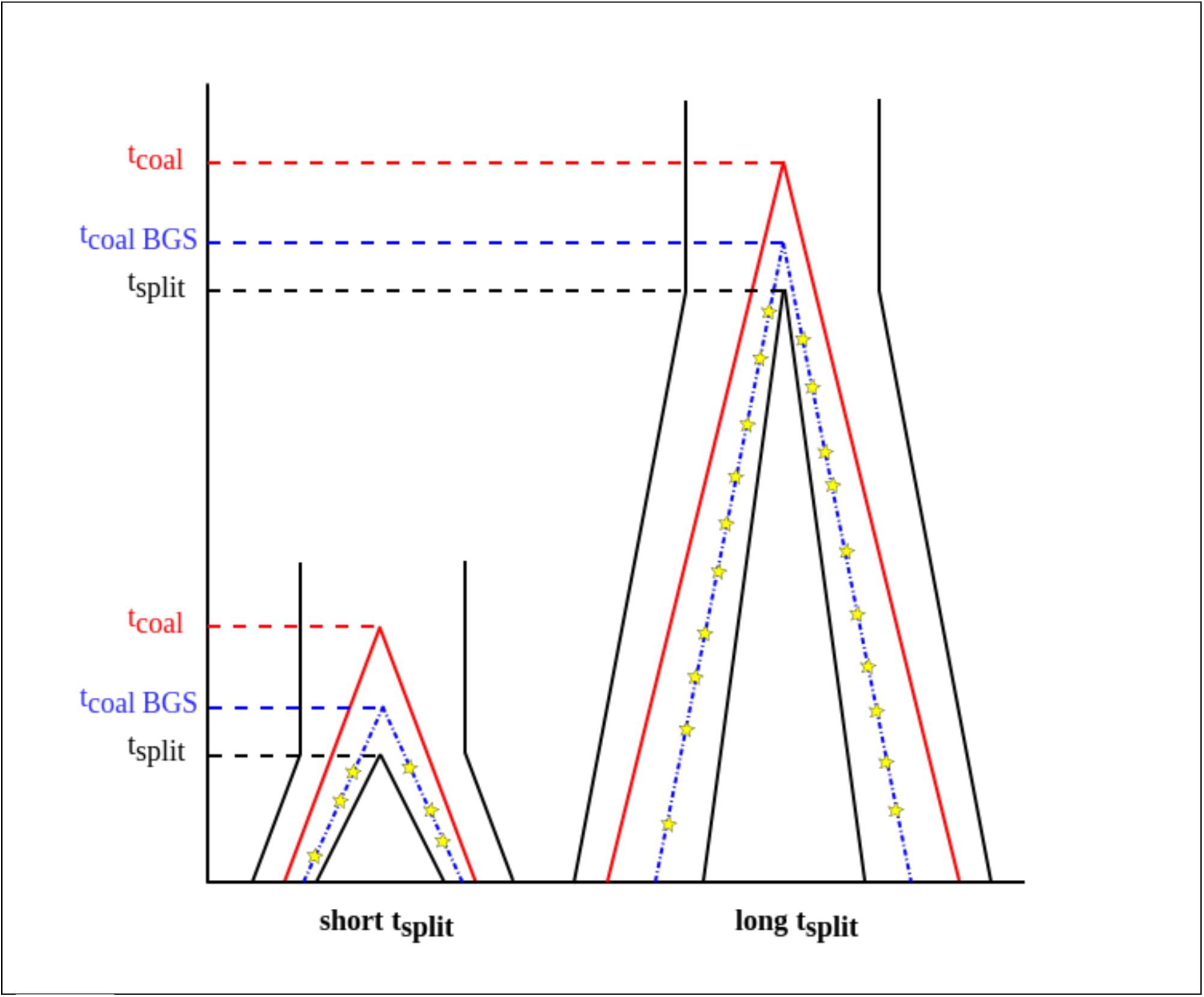
Models of how genealogies are affected by selection at linked neutral sites. The genealogies on the left represent species with a short split time such as human and chimpanzee. The genealogies on the right represent species with a long split time such as human and mouse. Red lines represent two lineages and their coalescent times. Blue lines represents two lineages and their coalescent time when there is selection at linked neutral sites in the ancestral population. Yellow stars denote mutations accumulating on each of the two lineages after they split. Note that with the longer split time, the proportion of the genealogy attributed to the ancestral population decreases.

In addition to conflicting conceptual predictions about the expected effect of selection on divergence at linked neutral sites, empirical studies have also been ambiguous. While some studies found no evidence for a correlation between divergence and recombination such as in Drosophila (Begun and Aquadro 1992; McGaugh et al. 2012) or in yeast (Noor 2008), other studies have reported correlations between divergence and recombination in Drosophila (Kulathinal et al. 2008; Begun et al. 2007). Further, positive correlations between human-chimpanzee divergence and human recombination rate (Hellmann et al. 2005; Cai et al. 2009; Lohmueller et al. 2011), human-macaque divergence and human female recombination rate (Tyekucheva et al. 2008), or human-baboon divergence and human recombination rate (Hellmann et al. 2003) have been reported. Intriguingly, several studies (Lercher and Hurst 2002; Mouse Genome Sequencing Consortium et al. 2002; Hardison et al. 2003) reported a positive correlation between human-mouse divergence and human recombination rate, but this empirical finding has not been interpreted in a model-based population genetic framework since its publication over 10 years ago. Finally, even though there was evidence for a strong reduction in human-chimpanzee divergence and human-macaque divergence surrounding genes (Tyekucheva et al. 2008; McVicker et al. 2009), McVicker et al. attributed the reductions seen for human-dog divergence to variation in mutation rates. Thus, the degree to which divergence is affected by selection across species with different split times remains elusive.

Determining whether and how selection affects linked neutral divergence is critical to understanding the evolutionary forces influencing genetic variation and mutational processes. If selection in the ancestral population only has a limited effect on divergence, it would suggest correlations between recombination and divergence to be evidence of mutagenic recombination. This may further suggest the need to consider recombination rates when modeling variation in mutation rates across the genome (Lercher and Hurst 2002; Hellmann et al. 2003; Pratto et al. 2014; Arbeithuber et al. 2015; Francioli et al. 2015). Because mutations rates have been difficult to reliable estimate in humans (Scally and Durbin 2012; Ségurel et al. 2014), understanding the biological factors influencing them will be of paramount concern for obtaining improved estimates. If, on the other hand, selection can affect linked neutral divergence, correlations between divergence and recombination or reductions of linked neutral divergence surrounding genes would suggest an abundance of selection affecting linked neutral sites (Sella et al. 2009). Selection affecting linked neutral diversity and divergence is at odds with the neutral and nearly neutral theories (Kimura 1983; Ohta 1973; Akashi et al. 2012), which have been the prevailing views in molecular population genetics for the last several decades. It would also suggest the need to consider the effects of selection when estimating mutations rates from neutral divergence.

Here we aim to examine the effects of selection on linked neutral divergence for species with a range of split times. We first present evidence that neutral divergence is reduced at sites linked to selected sites across a wide range of taxa, including those with split times as long as 75 million years ago. Many of these results cannot be caused by mutagenic recombination or biased gene conversion. Then, we present a theoretical argument as to how background selection can affect variation in neutral divergence across the genome, even for species with long split time such as human and mouse. We use simulations to explore the conditions under which background selection is predicted to affect linked neutral divergence and show that this effect is likely to be widespread across many taxa. Our empirical and simulation-based findings suggest the view that selection does not affect divergence at linked neutral sites between distantly diverged species needs to be revised.

## RESULTS

### Neutral divergence is reduced in regions of the genome with greater functional content

To understand the role of natural selection in reducing linked neutral divergence, we first examined the relationship between functional content and neutral divergence. We defined functional content as the proportion of sites within a window that overlapped with an exon or a phastCons region. We hypothesized that the effect of selection on linked neutral sites would be more pronounced at regions with greater functional content (Payseur and Nachman 2002). This hypothesis predicts a negative correlation between functional content and neutral divergence. To test this, we divided the human genome into non-overlapping windows of 100kb. To obtain putatively neutral divergence for each window, we filtered sites that overlapped with an exon transcript or a 44-way phastCons region (see the Methods section for further details). For human-mouse and human-rat divergence, we corrected for multiple mutations by applying the Kimura two-parameter model (Kimura 1980).

We found a negative correlation between functional content and neutral divergence between pairs of closely related species (Spearman’s ρ*_human-chimp_* = −0.3286, *P* < 10^−16^, Spearman’s ρ*_human-orang_* = −0.2985, **Figure 2A**, **Figure 2B**, **Supplementary Table 1**). Such a finding is consistent with the hypothesis that selection has reduced neutral divergence close to functional elements between closely related species. We next tested whether there was a correlation for more distantly related pairs of species, such as human-mouse and human-rat. These species were predicted to have diverged approximately 75 million years ago (Mouse Genome Sequencing Consortium et al. 2002) and, as such, current thinking would predict that there would be no correlation between divergence and functional content due to selection affecting linked neutral sites. Instead, we find that functional content is negatively correlated with neutral divergence, even between pairs of distantly related species (**Figure 2C**, **Figure 2D**). The magnitude of the correlation is greater than that seen for the closely related species (Spearman’s ρ*_human-mouse_* = −0.4885, *P* < 10^−16^, Spearman’s ρ*_human-rat_* = −0.4774, *P* < 10^−16^, **Supplementary Table 1**). Some features of the genome such as CpG sites or GC content are known to correlate with genic content (Kong et al. 2002; Hellmann et al. 2005; Cai et al. 2009; Lohmueller et al. 2011). To test whether these features confounded the correlations found in our data, we repeated our analyses removing potential CpG sites by omitting sites preceding a G or following a C (McVicker et al. 2009). The correlations were essentially unchanged after filtering CpG sites (**Supplementary Table 1**). We next computed partial correlations controlling for GC content. Similarly, we found that the correlations persisted (**Supplementary Table 1**) and were similar to or greater than that seen in the unfiltered data. These results suggest that the negative correlation between functional content and divergence is not driven by mutational properties associated with sequence composition.

**Figure 2:**
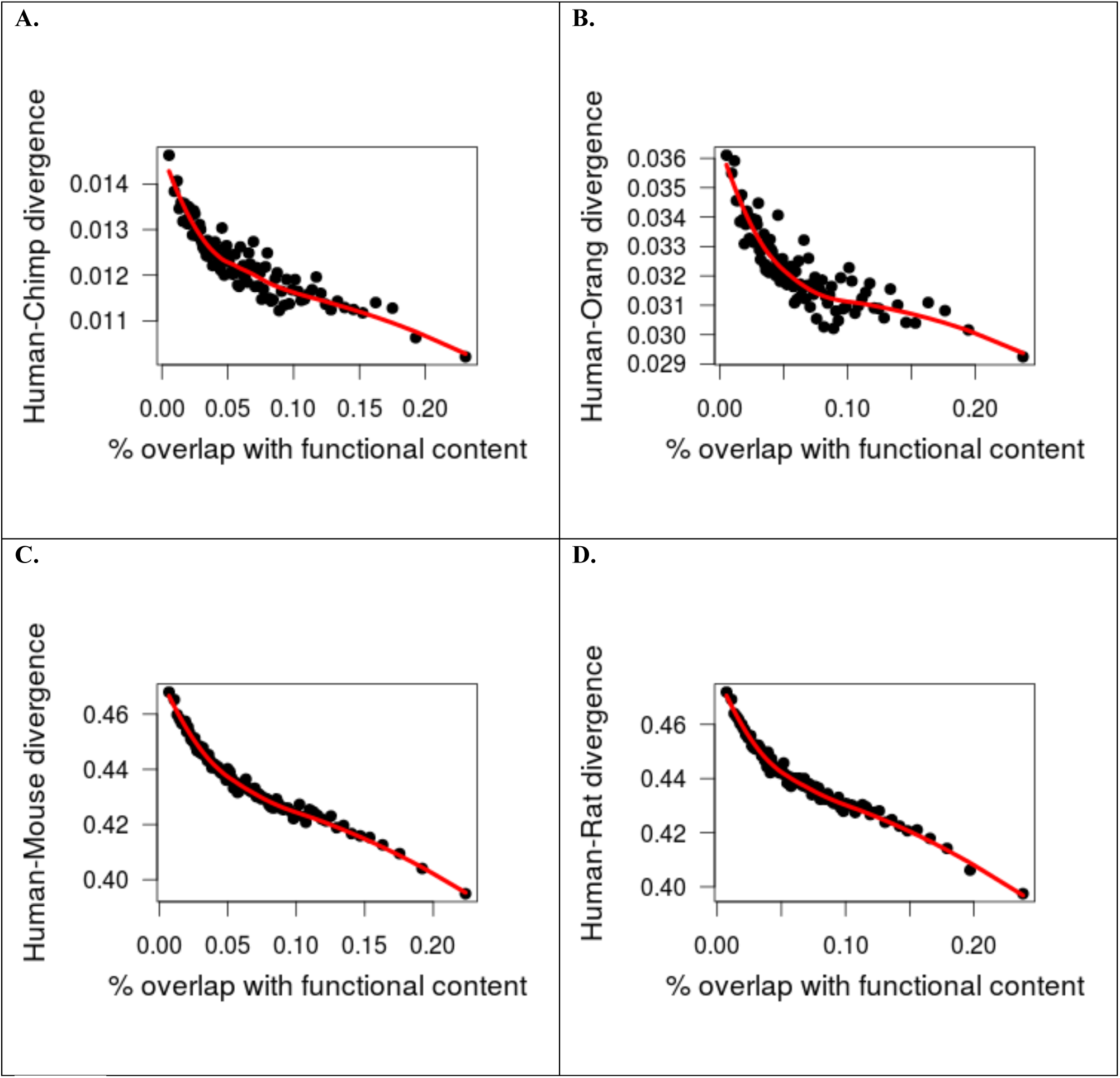
Neutral divergence is negative correlated with functional content. Each point represents the mean divergence and functional content in 1% of the 100kb windows binned by functional content. Red lines indicate the loess curves fit to divergence and functional content. Note that the last bin containing less than 1% of the windows was omitted from the plot. While the graph presents binned data, the correlations reported in the text are from the unbinned data. (A) Human-chimpanzee, (B) Human-orangutan, (C) Human-mouse, and (D) Human-rat divergence.

Biased gene conversion is an additional evolutionary force that has been shown to influence patterns of divergence (Galtier and Duret 2007; Duret and Arndt 2008). In this process, double-strand breaks in the DNA in individuals heterozygous for AT/GC variants will be preferentially repaired with the GC allele, resulting in AT ➔ GC substitutions occurring at a higher rate than GC ➔ AT substitutions (Duret and Arndt 2008; Duret and Galtier 2009; Berglund et al. 2009). To control for the effects of biased gene conversion on this analysis, we filtered out sites that could be affected. Since the exact sites affected by biased gene conversion are difficult to identify, we employed three different sets of filters, each with varying degrees of stringency. First, we filtered previously identified biased gene conversion hotspots (Capra et al. 2013). We obtained phastBias gBGC tracks for human and tabulated divergence excluding the sites that overlapped with the phastBias gBGC track. Negative correlations between functional content and divergence were essentially unchanged after filtering those sites (e.g. Spearman’s ρ*_human-mouse_* = −0.4838, **Supplementary Table 2**), suggesting that biased gene conversion clustered within the phastBias track cannot account for empirical correlations seen in the data.

However, since phastBias could only identify 25-50% of sites that are affected by biased gene conversion (Capra et al. 2013), we next filtered out any AT ➔ GC substitutions in regions of recombination hotspots, because it is thought that the effect of biased gene conversion is strongest in those regions (Glémin et al. 2015). To identify sites that fell into recombination hotspots, we used the double-strand break map (Pratto et al. 2014). Note, we did not wish to use recombination hotspots identified from patterns of linkage disequilibrium as the power to identify such hotpots may be related to SNP density, which may in turn be influenced by the degree of divergence between species. We filtered any substitutions where one species carried an A or T allele and the other carried a C or a G (i.e. weak to strong or strong to weak substitutions). Again, negative correlations between functional content and divergence were essentially unchanged after filtering those sites (e.g. Spearman’s ρ*_human-mouse_* = −0.4480, **Supplementary Figure 1**, **Supplementary Table 2**). Finally, we employed a more stringent filter for biased gene conversion by filtering any AT ➔ GC substitution genome-wide. Similar to what was seen with the other filters, the correlation between functional content persisted (Spearman’s ρ = −0.4710, **Supplementary Table 2**). In sum, our results suggest that the reduction in neutral divergence due to selection in the ancestral population has persisted to the present time. Importantly, because human-mouse divergence was computed only at putatively neutral sites, this pattern is unlikely to be driven by the direct effects of purifying selection removing deleterious mutations (see Discussion).

### Neutral human-rodent divergence correlates with background selection in primates

Next, we examined the relationship between human-mouse divergence and the strength of background selection across the genome inferred from divergence within primates (McVicker et al. 2009). This strength of background selection is captured by the *B-*value, which represents the degree to which neutral variation at a given position is reduced by selection relative to neutral expectations. While McVicker et al. found that divergence between primates was indeed reduced due to background selection, they did not consider human-mouse divergence in their analyses and did not model background selection within the human-dog ancestor. As such, there is no a priori reason why the *B-*values of McVicker et al. should be related to human-mouse divergence.

Nevertheless, we found a positive correlation between human-mouse divergence and the *B-*values from McVicker et al. (Spearman’s ρ = 0.5459, *P* < 10^−16^, **Figure 3A**, **Supplementary Table 3**). We also found a correlation between human-rat divergence and the *B-*values (Spearman’s ρ = 0.5242, *P* < 10^−16^, **Figure 3B**, **Supplementary Table 3**). These results suggest that regions of the genome that show stronger signals of background selection within primates (lower *B-*values) tend to show lower levels of human mouse divergence. Additionally, the positive correlation remained after filtering sites that may be within a CpG context or affected by gene conversion (Spearman’s ρ = 0.4404, **Supplementary Figure 2**, **Supplementary Table 4**), suggesting that some other process must be driving it. Our interpretation of this finding is that regions of the genome that are affected by background selection within primates had also been affected by background selection in the human-mouse ancestral population, and that this latter signal is still detectable in patterns of human-mouse divergence. Thus, this a second line of evidence suggesting that selection can affect linked neutral divergence.

**Figure 3:**
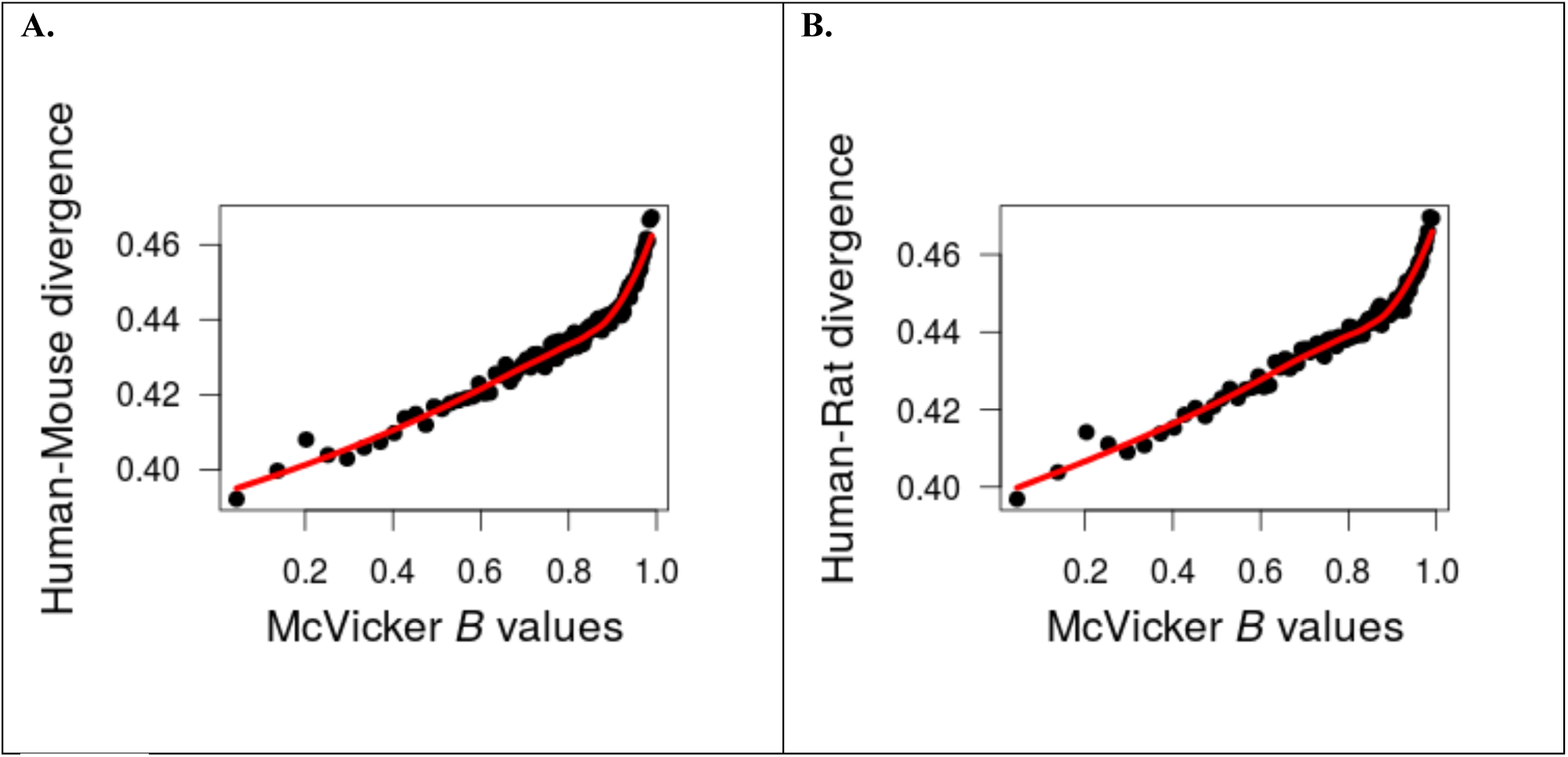
Neutral divergence is positively correlated with the strength of background selection inferred from primates (McVicker’s *B-*value). Each point represents the mean divergence and *B-*value in 1% of the 100kb windows binned by *B-*values. Red lines indicate the loess curves fit to divergence and recombination. Note that the last bin containing less than 1% of the windows was omitted from the plot. While the graph presents binned data, the correlations reported in the text are from the unbinned data. (A) Human-mouse and (B) Human-rat divergence.

### Divergence of closely and distantly related species is positively correlated with human recombination

Third, we examined the relationship between human recombination rates from the deCODE genetic map (Kong et al. 2010) and neutral divergence. We found a positive correlation between human recombination and neutral divergence between pairs of closely related species (Spearman’s ρ*_human-chimp_ =* 0.2564, *P* < 10^−16^, Spearman’s ρ*_human-orang_* = 0.2496, *P* < 10^−16^, **Figure 3A**, **Figure 3B**, **Supplementary Table 5**) as well as pairs of distantly related species (**Figure 3C**, **Figure 3D**). The magnitude of the correlation for distantly related species is about half that seen for the closely related species (Spearman’s ρ*_human-mouse_* = 0.1432, *P* < 10^−16^, Spearman’s ρ*_human-rat_* = 0.1337, *P* < 10^−16^, **Supplementary Table 5**).

To test whether natural selection could explain the correlation between genetic variation and recombination, we stratified windows of the genome into regions that are near genes and far from genes. We hypothesized that if natural selection reduced divergence at linked neutral sites, it would be more effective in removing divergence in regions overlapping genes because genes are more likely to be targets of natural selection (Lohmueller et al. 2011). Thus, we binned windows based upon the proportion that overlapped with a RefSeq transcript. We then computed Spearman’s ρ between divergence and recombination for each set of windows. We found that the correlation between divergence and recombination is stronger for windows that have greater overlap with RefSeq transcripts (**Supplementary Figure 3A**), suggesting that the correlation was stronger in regions near genes as opposed to far from genes. This result is consistent with the hypothesis that natural selection reduces divergence at linked neutral sites. After filtering for CpG sites or controlling for GC content, the correlations persisted (**Supplementary Table 5**) and were similar to or greater than that seen in the unfiltered data. These results suggest that the positive correlation between recombination and divergence is not driven by mutational properties associated with sequence composition.

Because biased gene conversion is thought to occur at a greater rate in regions of the genome with higher recombination rates, we were especially concerned that it could be driving the correlation between divergence and recombination. However, positive correlations between human recombination and divergence were essentially unchanged after filtering the sites within the phastBias track sites (e.g. Spearman’s ρ*_human-chimp_* = 0.249, **Supplementary Table 6**), After filtering AT➔GC sites within human double-strand break hotspots, the overall correlation between human-chimpanzee divergence and human recombination was slightly weaker, though still significant (Spearman’s ρ*_human-chimp_* = 0.2351, **Supplementary Table 6**), suggesting that biased gene conversion is unlikely to explain the correlation between human-chimpanzee divergence and human recombination. Though it remained formally significant, the overall correlation between human-mouse divergence and recombination substantially decreased after this filtering (Spearman’s ρ*_human-mouse_* = 0.0426, *P*<10^−5^, **Supplementary Table 6**). This suggests that we cannot exclude the possibility that biased gene conversion could be driving much of the genome-wide correlation between recombination and divergence when considering distantly related species. However, the presence of biased gene conversion does not negate the possibility that natural selection could also still affect divergence. Thus, we hypothesized that if natural selection could reduce neutral divergence at linked sites, it may still be detectable in windows near genes. Therefore, we stratified the windows based on the percentage of sites of each window that overlapped with a RefSeq transcript. The correlation between human-mouse divergence and human recombination was greater than 0.1 when considering windows with at least 10% overlap with a RefSeq transcript (**Supplementary Figure 3B**, **Supplementary Figure 4**). This implies that even after stringent filtering for biased gene conversion, genic windows still show a stronger correlation between recombination and divergence than do non-genic windows. Further, the fact that genic windows still show a significantly positive correlation between human-mouse divergence and recombination suggests that biased gene conversion cannot explain all of the patterns and that some other evolutionary force, such as selection at linked neutral sites, must be invoked.

**Figure 4:**
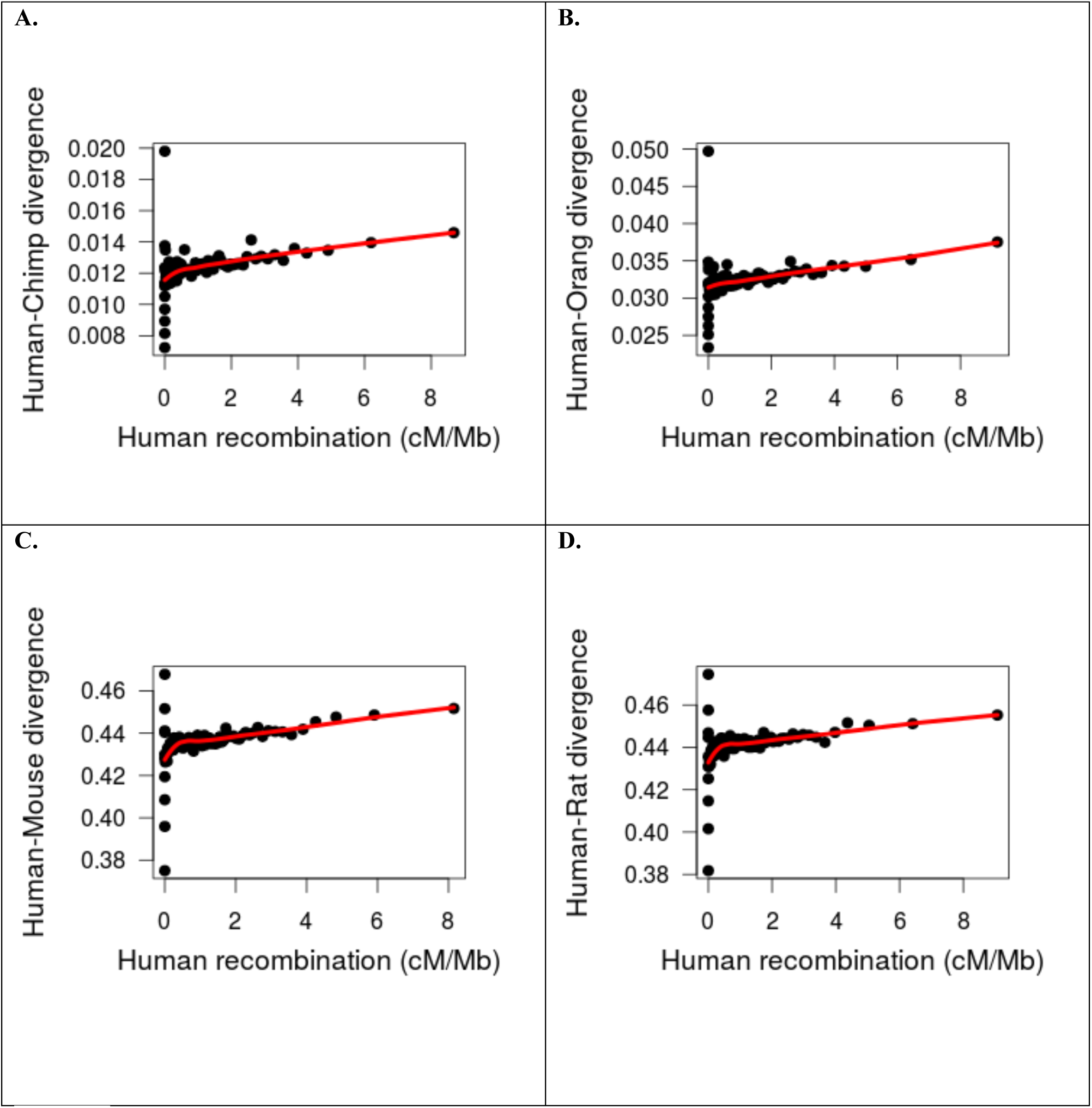
Neutral divergence is positively correlated with human recombination rate. Each point represents the mean divergence and recombination in 1% of the 100kb windows binned by recombination rate. Red lines indicate the loess curves fit to divergence and recombination. Note that the last bin containing less than 1% of the windows was omitted from the plot. While the graph presents binned data, the correlations reported in the text are from the unbinned data. (A) Human-chimpanzee, (B) Human-orangutan, (C) Human-mouse, and (D) Human-rat divergence.

After filtering all AT ➔ GC substitutions genome-wide, the correlation between human-chimpanzee divergence and human recombination was reduced, but remained significant (Spearman’s ρ = 0.1475, **Supplementary Table 6**). The correlation between human-mouse divergence and human recombination became negative after filtering all weak to strong and strong to weak substitutions (Spearman’s ρ*_human-mouse_* = −0.060, *P* < 10^−6^). However, the potential effect of background selection in reducing neutral divergence at linked sites was again evident when we stratified windows overlapping genes and those not overlapping genes. We found that the correlation between human-mouse divergence and human recombination is significantly positive (Spearman’s ρ*_human-mouse_* = 0.0985) at windows with >75% overlap with a RefSeq transcript (**Supplementary Table 7**).

In summary, filtering out sites that could be under the influence of biased gene conversion using different criteria suggested that biased gene conversion may be driving much of the genome-wide correlation between recombination and divergence at distantly related species. However, even after more stringent filtering, the correlation between recombination and divergence could still be detected when considering windows that are near genes. Therefore, some other mechanisms must be invoked to explain this pattern. Further, other patterns that are more stable through the course of evolution such as the proportion of sites that are functional and the amount of background selection in the ancestral primate populations suggest that selection has affected linked neutral divergence.

### Theoretical models predict a substantial contribution of background selection to variation in neutral divergence even for old split times

Any effect of background selection on the variation in neutral divergence across the genome can only result from its effect on divergence in the ancestral population, since deleterious mutations do not affect the fixation rate at linked neutral sites (Birky and Walsh 1988). An old split time between two species leads to only a small proportion of the total divergence having been accumulated in the ancestral population of the two species. As such, one would expect that for old split times, the impact of background selection on variation in divergence across the genome is negligible.

However, here we show that even when the amount of divergence that accumulated in the ancestral population (*d_a_*) is small relative to the total divergence (*d_t_*), there can still be a substantial amount of variation in divergence across the genome being influenced by background selection. We analyze a simple two-locus model to explore the influence of ancestral population size (*N_a_*), mutation rate (μ) and strength of background selection (*B*) on the proportion of variance in divergence that can be explained by background selection (**Figure 5A**). Recombination in the ancestral population leads to a distribution of coalescent times within each locus, with an average coalescent time of 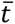. We assume that recombination rate within each locus is large enough, such that there is no variation in 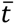 for a fixed value of B, i.e. 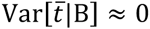. A difference in 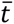 between loci is therefore only attributable to differences in background selection: 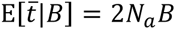. Further, variation in ancestral (*d_a_*) and total (*d_t_*) divergence results from a Poisson distributed number of mutations added to the genealogy, such that *Var* [*d_a_* |*B*] = [*d*_a_|*B*] = 4*N_a_BμL* and *Var*[*d_t_*|*B*] = *E*[*d_t_*|*B*] = *E[d_a_*|*B*] + 2*t_split_μL*. Here, *L* is the sequence length of a locus. The law of total variance can be used to compute the variance in total divergence across loci with varying levels of background selection:

**Figure 5:**
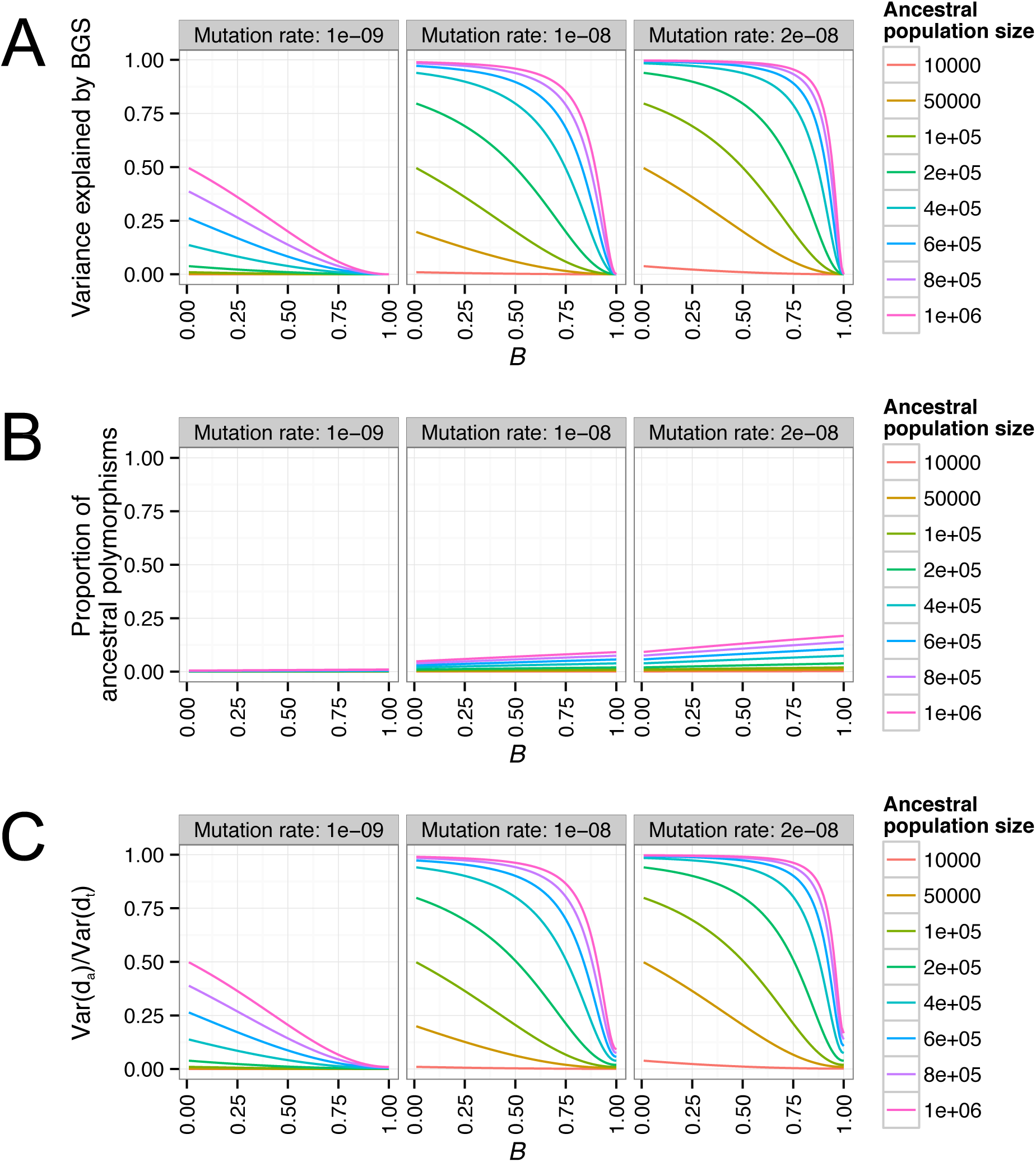
A two-locus model for the effect of background selection on divergence. (A) The variance in divergence between two loci explained by background selection as a function of the strength of background selection at the second locus (*B_2_*). (B) The expected proportion of divergence due to polymorphism in the ancestral population as a function of *B_2_.* (C) The variance in divergence between the two loci explained by polymorphism in the ancestral population as a function of *B_2_.* Different columns denote different mutation rates. Colored lines denote different ancestral population sizes (*N_a_*). Note that the variance in divergence attributable to background selection is greater than the expected proportion of divergence contributed by ancestral polymorphisms.

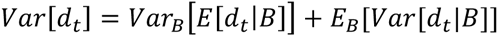

Thus, variance in total divergence can be decomposed into variance due to background selection and variance due to the mutational process. For simplicity, the first locus experiences no background selection (*B*_1_ *=* 1), and the second locus experiences some fixed amount of background selection (0 ≤ *B*_2_ ≤ 1). Under this model, we can compute the variance due to background selection as:

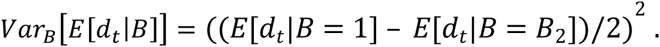

We then compute the variance due to the mutational process as:

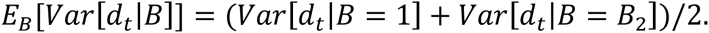

We assume an old split time, such that the divergence that accumulated from present time to population split is similar to the human-mouse divergence (40%). Both loci have a sequence length (*L*) of 100kbp. Our theoretical analysis of variance approach shows, that even with this old split time and a very small ancestral mutation rate of 1 × 10^−9^/bp, more than 20% of variation in divergence can be attributed to background selection in the ancestral population, as long as the ancestral population size is not too small (>600,000) and the effect of background selection is strong (*B_2_* < 0.2; **Figure 5A**). Note that under these conditions, the proportion of divergence that accumulated in the ancestral population can be as low as 0.3% (**Figure 5B**). However, the proportion of the variance in divergence that is attributable to the ancestral population is larger than 20% (**Figure 5C**), mainly due to background selection leading to differences in 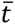 between loci. With a larger mutation rate (2 × 10^−8^/bp), one can observe a strong influence of background selection on variation in divergence even when ancestral population size is relatively small (>50,000). When assuming a moderately large population size of 200,000, and a moderate strength of background selection (*B_2_ =* 0.75), then as much as 50% of variance in divergence can be explained by background selection. Nonetheless, the proportion of divergence that accumulated in the ancestral population in this case is still only 3.4%. Our results demonstrate that, even for old split times where the vast majority of divergence accumulated after the population split, a moderate effect of background selection in the ancestral population can have a strong influence on the variation in divergence between genomic regions.

### Coalescent simulations predict background selection can reduce neutral divergence between species with long split times

Because the theoretical model described above assumes free recombination within windows and only considers a pair of loci at a time, we used coalescent simulations to examine whether background selection could generate this correlation under more realistic models. We modeled the effect of background selection as a reduction in the ancestral population size using the *B-*values estimated in McVicker et al. (see the Methods section for further details). For each window of the genome, we used a two-part strategy to simulate a neutral genealogy that was affected by background selection. The first part considered the genealogy in the ancestral population. This was done by simulating a neutral ancestral recombination graph (ARG) in a constant size population for a sample size of two using the population-scaled recombination rate *4N_a_Br,* where *N_a_* is the ancestral population size, and *r* is the recombination rate for the window, which we took from the deCODE genetic map. The ARG for the ancestral population used a population-scaled mutation rate *4NoBμ_a_,* where *μ_a_* is the ancestral mutation rate, which we set to 2.5 × 10^−8^ for these simulations. The second part of the genealogy considered the divergence that accumulated from time *t_split_* till the present day. Mutations were added to this part of the genealogy following a Poisson process (see Methods for more details).

We found that across all population sizes and split times examined, background selection generated a positive correlation between recombination and divergence as well as a positive correlation between divergence and *B-*values, even for pairs of species that split up to 100*N* generations ago (**Figure 6**). Interestingly, this correlation remained strong even when the proportion of the divergence due to ancestral polymorphism was small. For example, for a pair of populations with t_Split_=100*N* generations and an ancestral population of size 50,000, only 1.53% of the divergent sites are due to ancestral polymorphism. However, this model predicts a correlation of 0.2114 between recombination and divergence and a correlation of 0.377 between recombination and *B-*values. Although ancestral polymorphism only contributes in a small way to the total divergence, the variance in the amount of ancestral polymorphism across the windows accounts for nearly 60% of the variance in divergence across different windows (**Supplementary Figure 5**). In general, the correlations decreased as both the split time increased and the size of the ancestral population decreased (**Figure 6**). This behavior is expected as the contribution of the variance in levels of ancestral polymorphism to the variance in divergence decreases with increasing split time and decreasing ancestral population size (**Supplementary Figure 5**). These results are in agreement with the theoretical results presented for the two-locus models.

**Figure 6:**
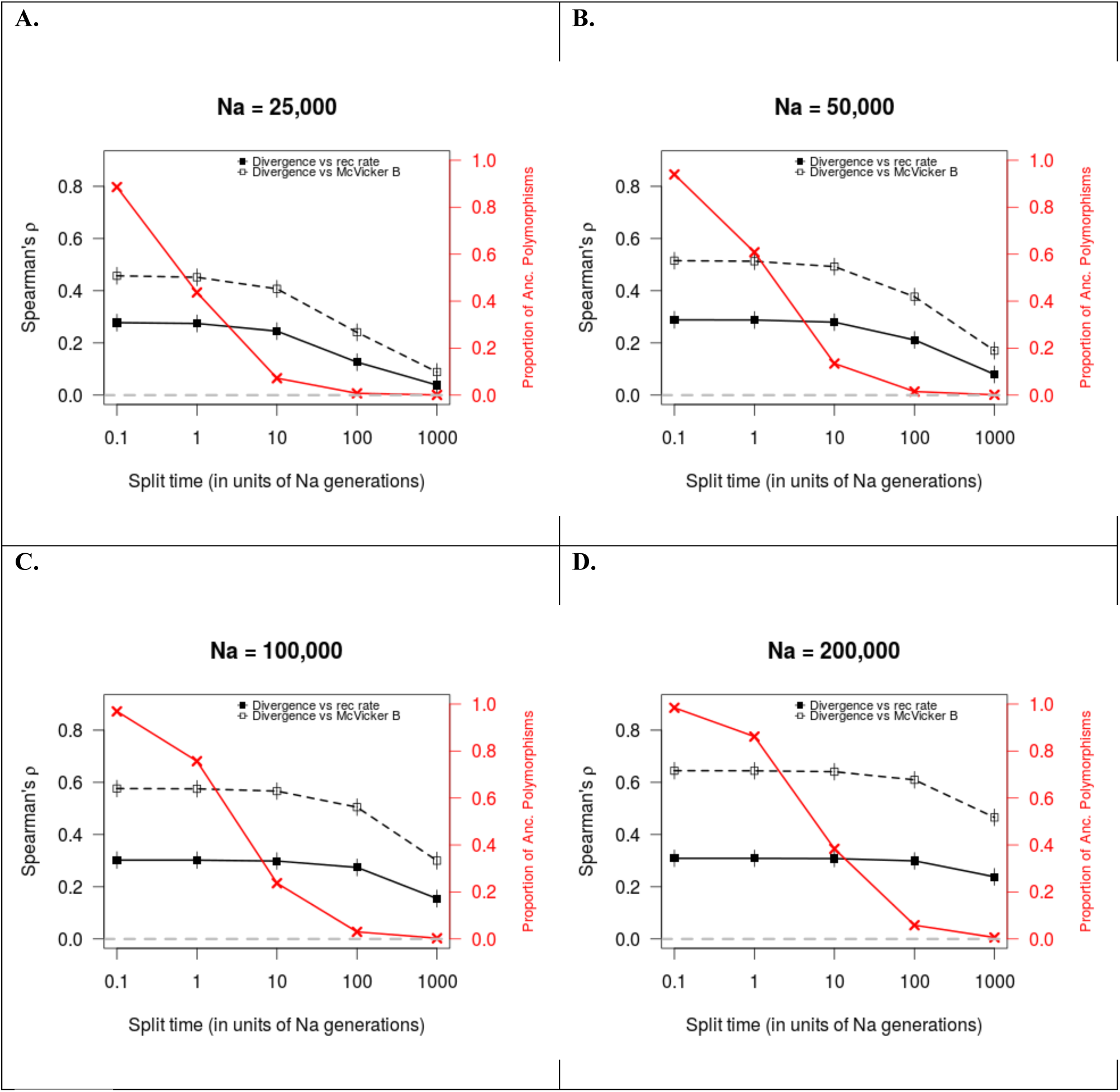
Background selection is predicted to affect neutral divergence across a range of split times and ancestral population sizes. Solid line shows the expected correlation coefficients (Spearman’s ρ) between neutral divergence and recombination rate as a function of split time. Dashed line shows the expected Spearman’s ρ between neutral divergence and McVicker’s *B-*values as a function of split time. Red lines denote the proportion of the divergence due to polymorphism that arose in the ancestral population. Error bars denote ± one standard error of the mean. Panels A-D denote difference in ancestral population sizes (*N_a_*). Note that the correlations are greater than 0 for a range of split times and ancestral population sizes, even when the proportion of divergence due to ancestral polymorphism is low.

We also explored the effect of background selection on divergence using additional sets of *B-*values. Specifically, we used the theoretical predictions of the effect of background selection as a function of *U,* (the deleterious mutation rate), the recombination rate, and *sh* (the heterozygous selection coefficient). We then picked values of *U* and *sh* that produced a distribution of *B-*values with mean of 0.8, the predicted mean strength of background selection in humans (McVicker et al. 2009), and 0.6, the mean predicted strength of background selection in *Drosophila* (Comeron 2014). For these *B-*values, we found that background selection still could generate a positive correlation between recombination and divergence with very ancient split times (**Supplementary Figure 6**). As before, we found that this pattern persisted across different ancestral population sizes (**Supplementary Figure 6**).

### Background selection in the human-mouse ancestor can explain genomic variation in neutral divergence

We tested whether background selection could explain our empirical observations regarding neutral human-mouse divergence. To do this, we used a coalescent simulation approach, which included background selection in a manner similar to that described above. Here we used parameter values that are more plausible for human-mouse populations. For example, we used an average mutation rate of μ*_a_* = 2 × 10^−8^ per generation for the human-mouse ancestral population. We included mutation rate variation across windows of the genome by drawing from a gamma distribution. Interestingly, a gamma distribution used for human mutation rates that had been calibrated by human-chimpanzee divergence (Voight et al. 2005) predicts a greater variance in human-mouse divergence than seen empirically. Thus, we selected the values of the parameters for a gamma distribution as well as the ancestral human-mouse population size to matched the mean and standard deviation of our observed human-mouse divergence (see Methods, **Supplementary Figure 7**). Ultimately, this model uses a human-mouse ancestral population size of 600,000. Given these parameters (**Supplementary Figure 8**), we then simulated 200 sets of genome-wide windows and computed Spearman’s correlation between divergence and the McVicker *B-*values for each replicate.

When considering models without background selection (i.e. *B*=1 for all windows), the average value of Spearman’s ρ was 0.025, and none of the 200 simulation replicates approached the value of Spearman’s ρ seen empirically (0.441, **Figure 7**). This result suggests that values of Spearmans’ ρ as large as that seen empirically cannot be generated by the neutral coalescent process. On the other hand, when modeling background selection using the McVicker *B-*values, the average Spearman’s ρ was 0.4461 which was comparable to the Spearman’s ρ computed from empirical human-mouse divergence (**Figure 7**). In sum, our results suggest that a model with background selection in the ancestral population can generate a positive correlation between the *B-*values and human-mouse divergence that is compatible to what is seen empirically, making background selection a plausible explanation for our empirical patterns.

**Figure 7:**
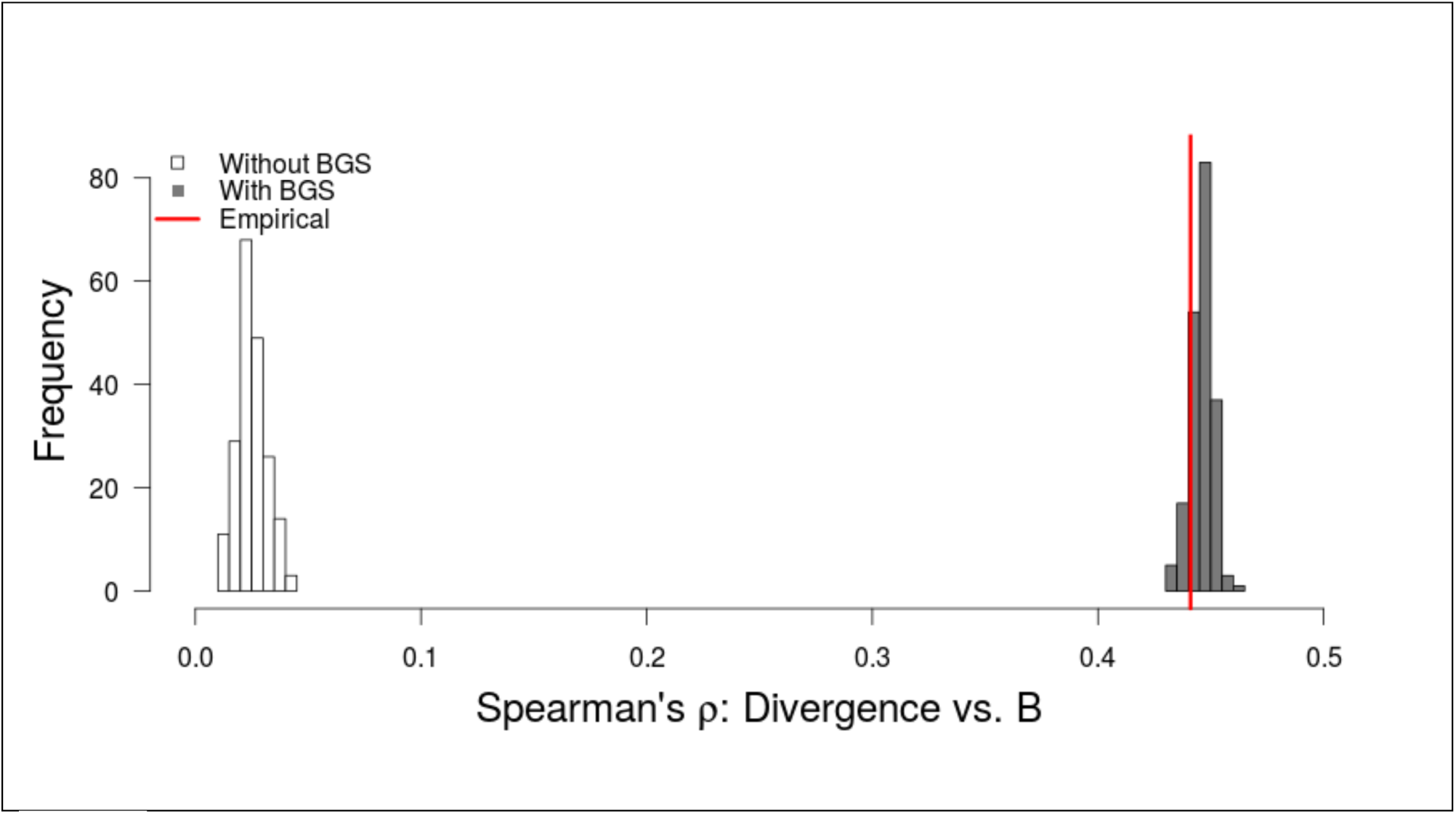
Models of background selection predict a correlation between neutral human-mouse divergence and McVicker’s *B-*values. Gray histogram denotes 200 simulations without including background selection. Blue histogram denotes 200 simulations incorporating background selection (see text). Red line represents the correlation computed from empirical human-mouse neutral divergence and McVicker’s *B-*values. Thus, plausible levels of background selection can match the observed correlation while neutral simulations cannot.

## DISCUSSION

Overall we show that neutral divergence, even between distantly related species, has been affected by natural selection. There are five lines of evidence in support of this assertion. The first line of evidence is that neutral divergence of distantly related species (i.e. human-mouse) negatively correlates with functional content. If natural selection is reducing neutral divergence at linked sites, the expectation is that neutral divergence would be lowest in regions with high functional content because natural selection tends to target those regions. Our empirical findings matched with the prediction (**Figure 2**). Second, human-mouse neutral divergence strongly correlates with the estimates of background selection in primates (**Figure 3**). These correlations persist after filtering out confounding factors, specifically CpG content and biased gene conversion sites. Third, we found a significant positive correlation between human recombination and putatively neutral divergence between closely and distantly related species (**Figure 4**). This correlation is evidence of selection affecting linked neutral sites because selection is predicted to reduce linked neutral divergence across a larger region of the genome in areas of low recombination than higher recombination. The fourth line of evidence is that the correlation between neutral divergence and recombination is stronger at regions that are near genes compared to regions that are far from genes (**Supplementary Figure 3** and **Supplementary Figure 4**). The reason why this is evidence of selection at linked sites is that natural selection is generally thought to affect regions of the genome that are functional, such as coding regions. Therefore, we would expect selection at linked sites to have more of an effect on regions near genes than regions far from genes. Our result is consistent with this prediction. Further, the overall correlation between recombination and neutral divergence of closely related species persists after filtering out sites that are potentially influenced by biased gene conversion. For distantly related species, even though the overall correlation between recombination and neutral divergence decreased, the signature of selection at linked sites was apparent when we stratified regions of the genome into whether they were near or far from genes. These results suggest that the signal of selection at linked sites when considering species with a long split time is quite subtle, perhaps weakened by changing recombination rates (see below), and that it could only be detected near genes because those regions ought to experience the strongest effect of selection at linked sites. The fifth line of evidence comes from our theoretical models and coalescent simulations. Our simulations show that even with very ancient split times (10*N* generations), when only a small amount (<10%) of the divergence comes from ancestral polymorphism, background selection is still expected to leave a signature on patterns of neutral divergence (**Figure 6**). Specifically, it can lead to a correlation between recombination and neutral divergence as well as a correlation between *B-*values and neutral divergence. These correlations occur because varying amounts of background selection across the genome can induce variation in the coalescent times across the genome, thus accounting for a large fraction of the variance in divergence across the genome.

While we initially observed s a genome-wide correlation between human-mouse divergence and recombination (**Figure 4**), it largely disappeared after filtering weak to strong and strong to weak substitutions that may have been affected by biased gene conversion. As such, much of this correlation seen in the data may be driven by biased gene conversion and not background selection. An obvious question is why we do not observe a correlation between human-mouse divergence and recombination rate genome-wide due to background selection when our simulations suggest one should be detected. One possibility could be that recombination rates may have changed throughout evolution. Even though human and chimpanzee do not share recombination hotspots, the broad scale recombination rates between human and chimpanzee tend to be conserved (Auton et al. 2012). Thus, the correlation between human-chimpanzee divergence and human recombination rate was still apparent. However, even the broad scale recombination rates differ between human and mouse (Jensen-Seaman et al. 2004). Changing recombination rates would likely erode the correlation between recombination and human-mouse divergence. Our simulations predict that if we had an estimate of the human-mouse ancestral recombination rate, then we would detect a positive correlation between human-mouse divergence and recombination rate. Nevertheless, the fact that in regions of the genome close to genes we could still detect a significant correlation between neutral human-mouse divergence and human recombination suggests that the signal from background selection is strong enough to have not been completely eroded by these changes in recombination rates.

These results suggest that examining the correlation between recombination and neutral divergence might not be the best measure to detect selection at linked sites. Perhaps other measures such as functional content or *B-*values that may not drastically change over the course of evolution are better alternatives. When we explored these other alterative patterns, we found correlations consistent with predictions from models of background selection. This result suggests that natural selection has affected neutral divergence and that the reason we do not see striking correlations between divergence and recombination rate may largely be due to a lack of power because of changing recombination rates.

Our coalescent simulations suggest that background selection can generate the correlation between human-mouse divergence and *B-*values seen in our data. However, these simulations make assumptions regarding the values of parameters like the ancestral human-mouse population size, generation times, and mutation rates over the last 75 million years. There is much uncertainty surrounding all of these parameters (Kumar and Subramanian 2002; Smith et al. 2002; Mouse Genome Sequencing Consortium et al. 2002; Hardison et al. 2003; Hodgkinson and Eyre-Walker 2011; Geraldes et al. 2011). For example, we assumed the ancestral human mouse population size was 600,000 and μ*_a_* = 2 × 10^−8^ per generation. This population size is the same order of magnitude as previously suggested for mice (Geraldes et al. 2011; Halligan et al. 2013). Assuming the human-mouse ancestor had a generation time of 1 year, our mutation rate would be higher than that previously suggested for mammals (Kumar and Subramanian 2002; Mouse Genome Sequencing Consortium et al. 2002; Hardison et al. 2003). However, a longer generation time would make the rates more comparable. We predict that μ*_a_* = 2 × 10^−9^ per generation, and 1 year per generation would require an ancestral population size of 4 million to generate the correlations seen in the empirical data. Nevertheless, the parameter space over which we expect background selection to generate a correlation between divergence and *B-*values is quite large, suggesting conclusions should not depend on the specific parameter values (**Figure 5**, **Figure 6** and **Supplementary Figure 6**).

While we found that models incorporating background selection predict correlations comparable to the empirical data, in principal, several other evolutionary processes may be able to generate these patterns. First, selective sweeps in the ancestral population could reduce divergence just like background selection. Given that we are unlikely to be able to survey patterns of polymorphism in the human-mouse ancestor in more than two lineages, it will be difficult or nearly impossible to distinguish between these two types of selection at linked neutral sites. Thus, one should interpret our use of *B-*values as reflecting a reduction in divergence due to the combined effects of both background selection and selective sweeps, as suggested in McVicker et al. (2009). A second possibility is that the negative correlation between divergence and functional content as well as the positive correlation between divergence and *B-*values could be driven by differences in mutation rate across the genome. Indeed, McVicker et al. attributed a positive correlation between *B-*values and human-dog divergence to the effects of variable mutation rates. However, for this mechanism to explain our results, it would require that mutation rates would have to be lower closer to genes and in regions of the genome thought to experience more background selection (i.e. in regions with lower *B-*values). Current evidence does not support either requirement. Recent estimates of the de novo mutation rate have not found any evidence of a reduction close to genes (Francioli et al. 2015). Further, Palamara et al. (2015) found that their estimates of the mutation rate do not differ as a function of *B-*values. Further, mutagenic recombination is unlikely to explain the empirical patterns in our study. Specifically, the correlation between divergence and functional content does not depend on recombination rate. The negative correlation between divergence and functional content remained strong when controlling for variation in recombination rates (**Supplementary Table 1**) suggesting our results are unlikely to be driven by mutagenic recombination. Nevertheless, our results do not rule out the possibility of mutagenic recombination and this topic certainly warrants further investigation. A final possibility is that the reduction in neutral divergence near genes and in regions with lower *B-*values could be due to the direct effects of purifying selection removing variation from the population. We attempted to mitigate this effect by considering putatively neutral sites that were not in exons or conserved across species. Thus, for the direct effects of purifying selection to explain our results, there would have to be many unidentified noncoding functional elements that are under purifying selection but are not detected by phastCons. Current studies provide, at best, limited support for such a scenario (Gulko et al. 2015).

Our finding that the genome-wide correlation between neutral human-mouse divergence and recombination rate substantially decreases when filtering weak to strong and strong to weak mutations suggests an important role for biased gene conversion at shaping patterns of divergence across genomes. Thus, our results are in line with previous reports of the importance of this effect (Duret and Arndt 2008; Duret and Galtier 2009; Capra et al. 2013; Berglund et al. 2009; Glémin et al. 2015).

One concern is whether poor alignment of divergent sequences influences our results. Specifically, we found that divergence at putatively neutral sites decreases with increasing functional content (**Figure 2**). It is possible that human and mouse sequences close to genes are easier to align because they have not diverged as much than sequences further from genes. Then, if the alignments further from genes contain more errors, there would appear to be more diverged away from genes. While this is conceptually possible, we do not believe it is driving the patterns seen in our analyses because poor alignments would not predict some of the patterns seen in our data. For example, if poorer alignment quality in regions far from genes was driving many of our results, then we would expect the correlation between recombination and human-mouse divergence to be strongest when considering all windows of the genome. When only considering genic windows, the correlation should decrease because we are only considering good-quality alignments. However, we observe the opposite pattern. The correlation between recombination and divergence actually increases with increasing the amount of genic sequence within a window. This result is more consistent with selection at linked neutral sites affecting divergence, rather than alignment quality.

Other studies have argued that background selection will not affect divergence between distantly related species because the genealogy in the ancestral population only comprises a small proportion of the total genealogy between one chromosomes from each of the two species (Birky and Walsh 1988; Hellmann et al. 2003; Begun et al. 2007; Cruickshank and Hahn 2014). This means that ancestral polymorphism will only account for a small proportion of the total divergence between distantly related species. It was thought that the signature of selection reducing the genealogy in the ancestral population would be diluted by the mutations that occurred since the split. As such, there would be no detectable signature of background selection. Our theoretical results and simulations show the proportion of ancestral polymorphism is actually a poor predictor of the correlation between divergence and recombination as well as between divergence and *B-*values. For example, consider a pair of species that split *N* generations ago with an ancestral population size of 25,000. In this model, 40% of the divergence is attributable to *ancestral* polymorphism (**Figure 6**). Now consider a second pair of species that split 100N generations ago where *N_a_*=200,000. Here <5% of the divergence is due to ancestral polymorphism. Previous intuition would suggest the effect of background selection would be stronger in the first pair of species because they split more recently and ancestral polymorphism makes a greater contribution to divergence. However, our simulations show the exact opposite pattern (**Figure 6**). The correlation between *B-*values and divergence is higher in the model with the more ancient split (Spearman’s *ρ* = 0.6098) than the one with the more recent split (Spearman’s *ρ* = 0.4517). Similar results are seen for the correlation between recombination and divergence. The reason for this discrepancy is that the main driver of these correlations is not the average amount of ancestral polymorphism, but rather the contribution of the variance in ancestral polymorphism to the variance in divergence. Even when ancestral polymorphism makes only a small contribution to the overall average divergence, a substantial amount of the variance in total divergence across the genome can still be explained by variance in ancestral polymorphism, particularly if the ancestral population size is large. Our theoretical results suggest that the variance in the amount of background selection in different regions of the genome can account for a lot of the variance in total divergence, even for species that split long ago. In sum, our theoretical results and simulations suggest that previous intuition has understated the importance of even small amounts of ancestral polymorphism on genome-wide patterns of divergence between species.

The importance of the ancestral population size at determining the effect of selection at linked neutral sites may explain a counter-intuitive pattern in our empirical analyses. We found that the negative correlation between functional content and human-rodent divergence was twice as large as that between functional content and human-primate divergence (**Supplementary Tables 1 and 2**). This pattern is counter-intuitive under previous thinking because ancestral polymorphism makes a greater contribution to human-primate divergence than human-rodent divergence. However, the human-rodent ancestor likely had a larger population than the primate ancestor. Our simulations suggest that in such cases, the species that diverged longer ago, where ancestral polymorphism makes a smaller contribution to divergence, can have stronger effects of background selection.

Our results have important implications for understanding patterns of genetic variation and divergence across genomes. First, our findings add to the growing literature suggesting the importance of background selection at shaping genome-wide patterns of variability across species (Wright and Andolfatto 2008; McVicker et al. 2009; Cutter and Choi 2010; Hernandez et al. 2011; Lohmueller et al. 2011; Charlesworth 2012a; Flowers et al. 2012; Hufford et al. 2012; Charlesworth 2012b; Cutter and Payseur 2013; Halligan et al. 2013; Campos et al. 2014; Comeron 2014; Slotte 2014; Wilson Sayres et al. 2014). Our new contribution to this literature is that natural selection affects divergence, even between distantly related species. Second, our findings indicate that correlations between recombination and divergence between distantly related species cannot, by themselves, be interpreted as evidence of mutagenic recombination. Our work suggests the need to consider whether models including background selection, as well as biased gene conversion, can explain the findings before attributing correlations to mutagenic recombination. Third, our work suggests that estimators of mutational properties that rely on contrasting patterns of divergence across different parts of the genome that may be differentially affected by background selection may yield biased results. This effect has been studied within primates in greater detail in recent work (Narang and Wilson Sayres 2015). Fourth, the fact that we detect evidence of background selection between distantly related species suggests that there is still some information about the distribution of coalescent genealogies across the genome. This distribution of coalescent genealogies can be exploited to obtain more reliable estimates regarding the human-mouse ancestral population size. While several methods exist to estimate ancestral demographic parameters from divergence (Takahata 1986; Rannala and Yang 2003; Wall 2003; Siepel 2009; Gronau et al. 2011), we suggest that these methods may be applicable for very distantly related species. Our findings that background selection can increase the variance in coalescent times across the genome suggest these methods as well as other statistical methods which seek to infer demographic history from the distribution of coalescent times across the genome, such as the PSMC approach (Li and Durbin 2011), should account for the increased variance in coalescent times across the genome due to background selection. Not accounting for background selection could result in inferring spurious demographic events to account for the additional variance in coalescent times across the genome. Lastly, our results suggest a need for caution when using patterns of divergence to calibrate neutral mutation rates. Some of the variation in divergence across the genome may be due to varying coalescent times—further accentuated by selection—rather than differing mutation rates (Gillespie and Langley 1979; Edwards and Beerli 2000). Future work could explore the extent to which selection at linked neutral sites can explain the discrepancies between different types of estimates of mutation rates (Scally and Durbin 2012; Ségurel et al. 2014).

## METHODS

### Correlation analyses

To calculate the divergence between each pair of species, we divided the human genome into 100kb non-overlapping windows. We obtained the pairwise (.axt) alignments between human/chimpanzee (hg18/panTro2), human/orang (hg18/ponAbe2), human/mouse (hg18/mm9), and human/rat (hg18/rn4) from the UCSC genome browser. These alignments are the net of the best human chained alignments for each region of the genome (Kent et al. 2003). Any bases that fell into to the regions of centromere, telomere and repeat were not considered further in this study. We also excluded sites that were missing in either of the species in the alignment. To obtain the putatively neutral regions, we filtered out sites that fell into coding and conserved regions. The coding regions were retrieved from the UCSC Table Browser with the following specifications: clade: Mammal, genome: Human, assembly: Mar. 2006 (NCBI36/hg18), group: Genes and Gene predictions, track: UCSC Genes, table: knownGene. Similarly, the conserved regions were retrieved from the UCSC Table Browser with the following specifications: clade: Mammal, genome: Human, assembly: Mar. 2006 (NCBI36/hg18), group: Comparative Genomics, track: Conservation, table: Vertebrate El (phastConsElements44way). For each window, we computed the total number of base pairs that passed the filtering criteria. To reduce variation, we only considered windows in which the total number of eligible base pairs was greater than 10,000. Then we computed the divergence by tabulating the number of base pairs that are different between the two species being compared. To account for multiple mutation hits for the distantly related species pairs (human-mouse and human-rat), we applied the Kimura two-parameter model (Kimura 1980). In addition, the recombination rate for each window was computed using the high-resolution pedigree-based genetic map assembled by deCODE (Kong et al. 2010). We then calculated Spearman’s ρ between divergence and recombination using the *cor* function in R.

### Filtering criteria

To filter out possible CpG sites, we excluded sites that were preceded by a C or were followed by a G in hg18 (McVicker et al. 2009). To control for the effects of biased gene conversion, we employed three different filters. First, we removed bases that overlapped with the phastBias track for humans (Capra et al. 2013). The phastBias track was retrieved from the UCSC Table Browser with the following specifications: clade: Mammal, genome: Human, assembly: Mar. 2006 (NCBI36/hg18), group: All Tables, database: hg18, table: phastBias Tracts 3. Second, confounding effects due to biased gene conversion were also controlled for by omitting AT ➔ GC substitutions in double-strand break regions. The double-strand break map obtained from Pratto et al. (2014) was used as a proxy for recombination hotspots. Rather than polarizing the directionality of the substitution, which may be error prone, we removed all divergent sites where one species had an A or a T nucleotide and the other had either a C or a G nucleotide. Thus, we only retained weak to weak or strong to strong mutations. Third, we removed all AT➔GC substitutions across the genome. This was done using the same steps as previously described.

### Simulations

To determine whether background selection could affect genetic divergence, we performed coalescent simulations that included background selection. We modeled background selection as a simple reduction in effective population size in the ancestral population (Charlesworth et al. 1993; Hudson and Kaplan 1995). This was done by scaling the ancestral population size *N_a_,* by the *B-*values. Unless otherwise noted, we used the *B-*values from McVicker et al. (2009). Each simulation replicate consisted of two parts. The first part modeled genetic variation in the ancestral population, and would include the effects of background selection. For each window *i,* we simulated an ancestral recombination graph (ARG) with a population-scaled recombination rate *4N_a_B_i_r_i_,* where *N_a_* is the ancestral population size, *B_i_* is the strength of background selection affecting window *i*, and *r_i_* is the recombination rate for window *i*. Mutations to the genealogy assuming a population-scaled mutations rate *θ*=4*N_a_*B*_i_*μ_a_,*_i_L_i_*, where μ_a_*_i_* is the ancestral per-base pair mutation rate for window *i* and *L_i_* is the number of successfully aligned neutral bases in window *i*. Simulations were done using the program ms (Hudson 2002). Note, we included recombination in the ancestral population because it affects the variance in coalescent times across windows and this variance in coalescent times will in turn affect the variance in levels of divergence, which will ultimately affect the strength of the correlation between divergence and recombination. Thus, we aimed to capture the appropriate variance to the extent possible. This part of the simulation generated the amount of divergence due to ancestral polymorphism, which we call *da*.

We then added the mutations that arose since (i.e. more recently) the split. The divergence from the present time to split time follows a Poisson distribution where the rate parameter equals the expected divergence between two populations. For the simulations in **Figure 6**, *d_s_ =* 2*t_split_μL* where *d_s_* is the expected divergence from the present time to the split time in the divergence model, *t_split_* is the split time, μ is the mutation rate, and *L* is the length of each sequence. When computing both *d_a_* and *d_s_* for the results in **Figure 6**, we assumed μ = 2.5 × 10^−8^ per base-pair per generation. For each window of the genome, *d_s_* was simulated using the *rpois* function in R. Finally, the total divergence within a window is the sum of divergence generated in the ancestral population (*d_a_*) and the divergence generated since the two species split (*d_s_*).

Due to the differences in generation times and mutation rates between the human and mouse lineages, we modified our approach for these simulations (**Figure 7** and **Supplementary Figure 8**). First, here *d_s_ =*(*t_mouse_ μ_mouse_+ t_human_μ_human_*)*L,* where *t_mouse_* is the number of generations on the lineage leading to the mouse from *t_split_* till the present day, *t_human_* is the number of generations on the lineage leading to human experienced from *t_split_* till the present day, μ*_mouse_* is the mutation rate along the mouse lineage, and μ*_human_* is the mutation rate along the human lineage. There is much uncertainty surrounding these parameters. However, the following values are broadly consistent with what has been reported previously and match the observed mean and standard deviation of human-mouse divergence (**Supplementary Figure 7**). First, we assumed *t_split_* = 75 million years ago. We then assumed mice have 1 generation per year, giving *t_mouse_* = 75 × 10^6^ generations. We assumed humans have 25 years per generation, making *t_human_* = 3 × 10^6^. We then set *μ_mouse_* = 3.8 × 10^−9^ per generation and *μ_human_* =3.75 × 10^−8^ per generation. These estimates are broadly consistent with previous reports and allow for approximately twice as much divergence on the mouse lineage as compared to the human lineage (Mouse Genome Sequencing Consortium et al. 2002). Thus, the expected value for *d_s_* was 0.3975 per site.

We assumed that μ*_a_* was equal to 2 × 10^−8^ per generation, which is the average of μ*_human_* and μ*_mouse_*. We accounted for variation in mutation rates across different regions of the genome by drawing μ*_a_* from a gamma distribution (Voight et al. 2005). We kept the ratio of μ*_a_* to μ*_mouse_* constant across all windows of the genome. For example, μ*_a_* / μ*_mouse_* = 5.26. Then if μ*_a_*_,i_ is the rate for the *i*^th^ region drawn from the gamma distribution, we set μ*_mouse_,_i_* equal to μ*_a_*_,i_ / 5.26. A similar procedure was used to find μ*_human,i_*. Increasing the variance in the mutation rate across regions increased the variance in divergence across windows of the genome and decreased the correlation between divergence and the *B-*values. We then examined different values of *N_a_* and parameters of the gamma distribution that matched the observed mean and standard deviation of the distribution of human-mouse divergence. We set *N_a_* = 600,000 and the shape parameter equal to 1212 and the scale equal to 1.65 × 10^−11^, which matched the observed mean and standard deviation of human-mouse divergence reasonably well (**Supplementary Figure 7**).

To explore alternate sets of *B-*values, we computed *B-*values using the following approach. We began with the theoretical prediction from (Hudson and Kaplan 1995) that 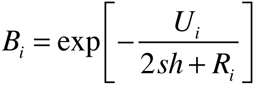. Note that *R_i_* represents the recombination rate for the *i^th^* window of the genome that is taken from the deCODE map. We then searched over values of *U* and *sh* that, in combination with the genome-wide distribution of *R_i_,* yielded mean values of *B* that were equal to 0.8 and 0.6. For *B* = 0.8, we used *U* = 0.00152 and *sh* = 0.003. For *B* = 0.6, we used *U* = 0.0054 and *sh* = 0.005.

## ACKNOWLEDGMENTS

We thank Melissa Wilson Sayres for helpful discussions. This research was funded by a UCLA Hellman Faculty Fellowship and an Alfred P. Sloan Research Fellowship in Computational & Molecular Biology to KEL. TNP was supported by the National Institutes of Health, under Ruth L. Kirschstein National Research Service Award (T32-GM008185).

